# Pheno-morphological screening and acoustic sorting of 3D multicellular aggregates using drop millifluidics

**DOI:** 10.1101/2024.07.22.604529

**Authors:** Leon Rembotte, Thomas Beneyton, Lionel Buisson, Amaury Badon, Adeline Boyreau, Camille Douillet, Loic Hermant, Anirban Jana, Pierre Nassoy, Jean-Christophe Baret

## Abstract

Three-dimensional multicellular aggregates like organoids and spheroids have become essential tools to study the biological mechanisms involved in the progression of diseases. In cancer research, they are now widely used as in vitro models for drug testing. However, their analysis still relies on tedious manual procedures, which hinders their routine use in large-scale biological assays. Here, we introduce a novel drop millifluidic approach to screen and sort large populations containing over one thousand multicellular aggregates. Our system utilizes real-time image processing to detect pheno-morphological traits in cellular aggregates. They are then encapsulated in millimetric drops, actuated on-demand using the acoustic radiation force. We demonstrate the performance of our system by sorting spheroids with uniform sizes from a heterogeneous population, and by isolating organoids from spheroids with different phenotypes. We anticipate that this work offers the potential to standardize drug testing on multicellular aggregates, which promises accelerated progress in biomedical research.

## Introduction

Over the past decade, three-dimensional (3D) multicellular aggregates (MCAs) have emerged as the new gold standard to investigate fundamental cell biology processes with a higher degree of physiological relevance compared with two-dimensional (2D) cultures (*1*). In particular, key phenomena like gene expression (*2*), cell-cell and cell-matrix interactions (*3*), physiology (*4*) and differentiation (*5*) have been shown to be better recapitulated in 3D models. The versatility of 3D models extends to a wide range of applications. In cancer research, multicellular spheroids composed of cancerous cells serve as in-vitro tumor models for disease modelling and drug testing (*6*). In cell therapy and regenerative medicine, organoids derived from patient cells or induced pluripotent stem cells (iPSCs) are used as building blocks to repair tissues (*7*). Recently, organoids have even been validated as an alternative to animal testing as stated in the US Food and Drug Administration Modernization Act 2.0 (*8*). This ever-growing interest for 3D cell models in biological studies pushes for the development of standardized methods to produce, analyze and screen them. The prerequisite for the production of 3D cell models is cell self-organization in a controlled environment (*9*). Traditional production methods like the hanging drop (*10*) and the spinning culture (*11*) have proven efficient for the formation of MCAs, but they demand considerable time investment. This makes them incompatible with experiments involving hundreds of samples. They also inherently introduce variability in the initial cell densities and nutrient concentrations, which subsequently gives rise to considerable sample heterogeneity (*12*). In contrast, microfluidic approaches enable high throughput production of MCAs and offer the possibility to control the chemical and mechanical properties of their microenvironment (*13–15*). Among others, the Cellular Capsules Technology (CCT) was designed to encapsulate cells in hollow alginate shells, allowing them to self-assemble into MCAs and grow in confined, niche-like microenvironments, hence producing thousands of MCAs per second (*16*). In this context, while most efforts were initially devoted to the production of MCAs, the bottleneck in the field of 3D biology has now shifted towards automatized, high-throughput characterization and manipulation methods.

The analysis of MCAs most often relies on user-dependent, manual methods. The key challenges result from their 3D nature and their wide range of sizes, from 50 *μ*m to 5 mm in diameter (*17*). A widespread approach consists in the dissociation of MCAs to perform single-cell analysis, or even in the lysis of the cells to perform biochemical assays (*18, 19*). These methods are however highly destructive and come with a complete loss of structural information. To analyze MCAs while preserving their integrity, optical microscopy remains the most adapted tool, since it allows to study MCAs at the multicellular level with a sub-cellular resolution (*20*). For instance, the growth dynamics and 3D internal organization of MCAs were unraveled thanks to advances in depth-resolved fluorescence microscopy (*21*), and in multi-plane image segmentation algorithms (*22*). Due to their complexity, these high-content approaches may only be applied to a small number of MCAs at a time. Combined developments in microfluidics and light-sheet fluorescence microscopy recently provided the possibility to analyze numerous MCAs in micro-engineered wells (*23*), or in manually-controlled flow (*24*). However, none of these approaches allows to manipulate MCAs for sorting purposes. Using MCAs in drug testing requires rapid screening of large populations of MCAs to gather statistically relevant information, and to isolate MCAs of interest for further analysis. To date, this would only be feasible through tedious pipetting of samples and time-consuming image acquisition.

Inspired by the development of flow cytometry in the field of single-cell analysis, we sought to develop a flow-based approach to address the pressing need for automated manipulation of MCAs. Since its invention in 1965 (*25*), flow cytometry has been massively adopted in biology facilities, especially through the advent of Fluorescence Activated Cell Sorting (FACS) for single-cell analysis (*26*). In flow cytometry, a suspension of cells continuously flows through a capillary where parameters of interest are measured in individual cells. They are then encapsulated into liquid drops by passing through a vibrating nozzle. Finally, the drops are deflected on-demand by submitting them to an electric field while they fall. However, the adaptation of flow cytometry to the analysis of MCAs has hardly been explored, mainly because of the clogging risks that arise when working with such large, weakly deformable objects. To our knowledge, only one group reported in 1987 the modification of a commercial flow cytometer to sort spheroids by increasing the size of the exit nozzle, hence allowing to study MCAs smaller than 100 *μ*m in diameter (*27*). Despite its pioneering nature, this approach was limited by the small size of the MCAs it could sort and its compatibility solely with detection techniques specific to standard FACS.

In addition to the challenges related to fluidics, scaling up from single cells to whole MCAs requires to redefine the parameters of interest that need to be measured. A classical analysis based on fluorescence intensity or light scattering cannot satisfactorily be used to characterize thick 3D objects. It would imply measuring averaged parameters over the whole MCAs, which comes with a loss of structural information. On the contrary, forming an optical image of a MCA gathers information across a whole surface, which yields a more complete description of their spatial organization. Such image-based approaches have only recently been unlocked for single cell analysis with the advent of microfluidic devices (*28, 29*). Not only microfluidic systems are suitable for imaging biological systems, but they are also compatible with many actuation methods previously implemented for single cell sorting: electrophoresis (*30*), dielectrophoresis (*31*), acoustophoresis (*32*), optical manipulation (*33*), or mechanical actuation (*34*). Recent developments in drop-based microfluidics further increased the versatility and the throughput of cell sorting by miniaturizing the principle of FACS into micrometric channels (*35*). Again, these approaches cannot be adapted for MCAs simply by increasing the channel dimensions. The large size of MCAs leads to a more predominant role of inertial forces, together with increased risks of sedimentation and clogging compared to single cells.

To address the limitations that currently prevent the widespread use of MCAs, we introduce a novel drop-based approach to perform Image-based Organoid Cytometry and Acoustic Sorting (ImOCAS). Like classical flow cytometers and droplet microfluidic devices, it carries out three primary operations: detection of a feature of interest in MCAs in flow, encapsulation of individual MCAs in liquid drops, and actuation of the drops of interest. Here, these steps were redesigned to allow the manipulation of biological samples up to several hundred microns in diameter. In ImOCAS, spheroids and organoids are continuously flowed through a square glass capillary where their morphological and phenotypical signatures are characterized on-the-fly using bright-field microscopy image analysis. They are then individually encapsulated in millimetric drops of culture medium which are sorted on-demand using the Acoustic Radiation Force (ARF) generated by a standing-wave acoustic field.

To illustrate the capabilities of ImOCAS, we screen large populations exceeding one thousand MCAs to extract statistical distributions of morphological and phenotypical features. We demonstrate its ability to accurately select spheroids of the same size from heterogeneous populations, which is a necessary initial step for subsequent drug testing. We also show its capacity to classify and separate plain MCAs from those containing a hollow core (lumen), which is a complex and time-consuming task when performed manually. The versatility, simplicity and generality of ImOCAS suggest its widespread adoption in 3D biology laboratories, with the potential to accelerate drug discovery and fundamental research thanks to high-throughput morphological and phenotypical screening of spheroids and organoids.

## Results

### Operating principle of ImOCAS

We first briefly explain the working principle of ImOCAS before detailing the challenges to overcome at each step in the following sections. *Figure 1* shows a schematic view of our system, which comprises three modules for the detection (i), encapsulation (ii) and actuation (iii) of MCAs.

**Figure 1.**
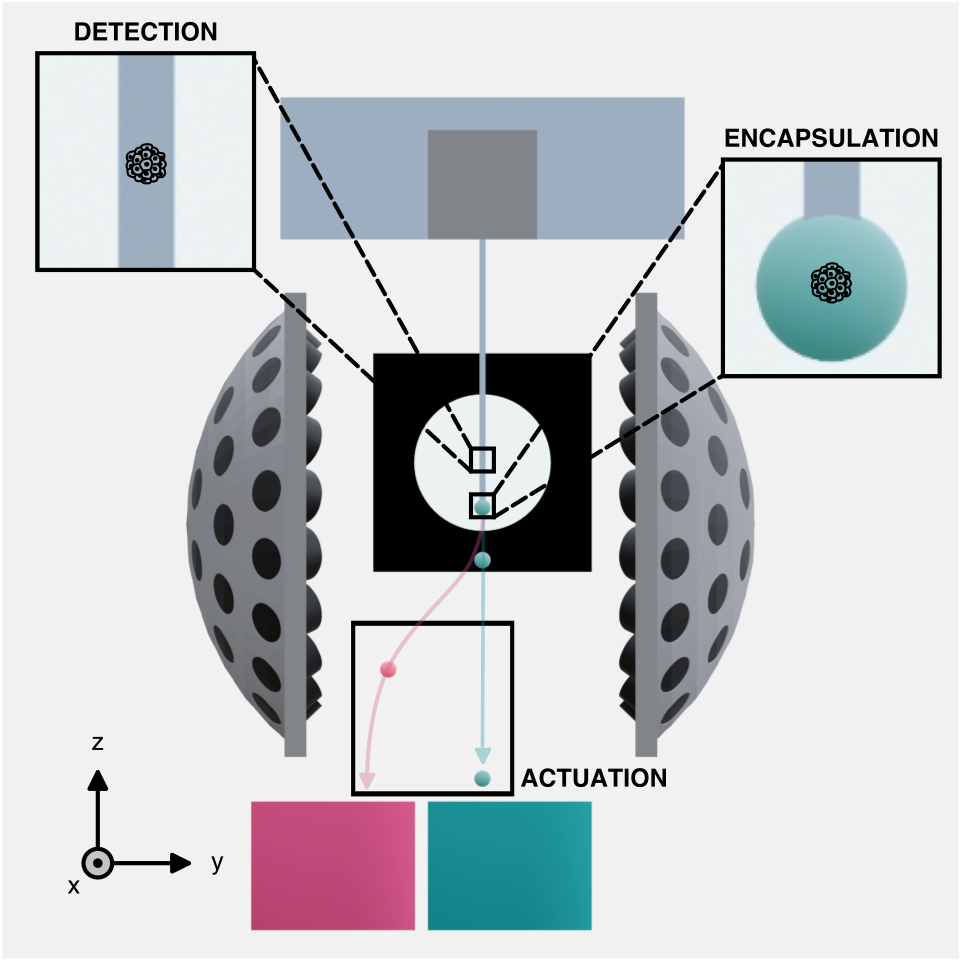
Working principle of ImOCAS: Detection, Encapsulation and Actuation. Using a simple microfluidic chip, MCAs are flowed into a glass capillary where a bright-field microscopy image is acquired. First, real-time image processing is used to characterize the morphology and the phenotype of MCAs, allowing to identify targets (detection). Then, each MCA is individually encapsulated in a drop of culture medium at a rate of ∼ 1 MCA per second (encapsulation). Finally, when a target MCA has been detected and encapsulated, the corresponding drop is acoustically deflected upon the activation of two spherical arrays of ultrasonic transducers (actuation). Axis arrows: 1.7 cm. Each element is at scale, colored arrows represent fictitious trajectories for visualization.

#### (i) Opto-fluidic detection

Real-time imaging and analysis of MCAs is required to characterize them with human-interpretable features. MCAs are dispersed in a culture medium solution and flowed through a square glass capillary. Glass capillaries are preferred to cylindrical capillaries to avoid optical aberrations. Micrographs are taken and analyzed on the fly at ∼ 100 frames per second (fps) to measure their morphological and phenotypical features, which are then compared to user-defined criteria to make a sorting decision.

#### (ii) Drop encapsulation

Each MCA is encapsulated in a millimetric drop of culture medium at the capillary exit. As detailed in a following section, the concentration of MCAs is optimized to encapsulate only one per drop, and to prevent the clogging of the exit capillary.

#### (iii) Acoustic actuation

The goal is to deflect individual drops using the ARF without altering the flow. A standing-wave acoustic field is generated by two arrays of ultrasonic transducers operating at 40 kHz. The arrays consist of two spherical caps, with their focal point placed close to the capillary exit. Upon detection of a target MCA, a standing-wave acoustic field is activated in proximity of the capillary exit to deflect the next drop towards a collection vial, while non-deflected drops are collected separately.

### Image-based MCA screening with on-the-fly detection of pheno-morphological features

Flow cytometry requires the measurement of a well identified set of parameters in a large number of objects, both for the collection of statistically relevant data and for the identification of potentially rare events. Here, we specifically aim at measuring morphological and phenotypical attributes of MCAs. More specifically, we analyze their sizes and shapes as first order discriminating parameters, and we monitor the presence or the absence of a lumen, which is a phenotypic property of epithelial tissues. We introduce a pipeline for rapid image-based characterization of MCAs, comprising: (i) the continuous acquisition of bright-field, monochromatic images upstream from the glass capillary exit, (ii) the detection of an MCA via binary image processing, (iii) the quantification of its morphological and phenotypical features, and (iv) its classification as target or waste based on user-defined criteria. Among the variety of shapes and topologies found across multicellular aggregates, we focus on two models: spheroids (plain ellipsoidal aggregates) of immortalized human embryonic kidney cells (HEK293T), very often encountered in tumor models, and cysts (spherical monolayers of epithelial cells surrounding a lumen) of induced pluripotent stem cells (iPSCs), from which organoids are often derived. Both spheroids and cysts are formed in hollow hydrogel shells using the CCT technique (see *Materials and Methods*).

*Figure 2A* summarizes the image processing steps performed in ImOCAS. Intensity thresholds and binary operations are applied to each bright-field image and allow to define four regions: *Outer, Inner, Border*, and *Centroid*. The projected area and the perimeter of the analyzed MCA is then simply derived by counting the number of pixels (px) in the *Outer* + *Inner* and Border domains, respectively. Note that splitting the object into its*Outer* and*Inner* regions is only required for discriminating between spheroids and cysts, where one has to take into account the presence of a lumen. Simple area detection is sufficient if only the morphological characterization of the MCAs is required. Our approach is compatible with on-the-fly analysis of ∼ 100 fps in 256 × 256 px^2^ images (*Fig. 2B*). The total processing times scale linearly with the total number of pixels in the image, which means the analysis throughput and/or the image size may be increased at will using a more powerful computer, if required. More complex analysis including shape fitting may also be implemented, taking into account that it slows the processing speed down (*Fig. 2B*).

**Figure 2.**
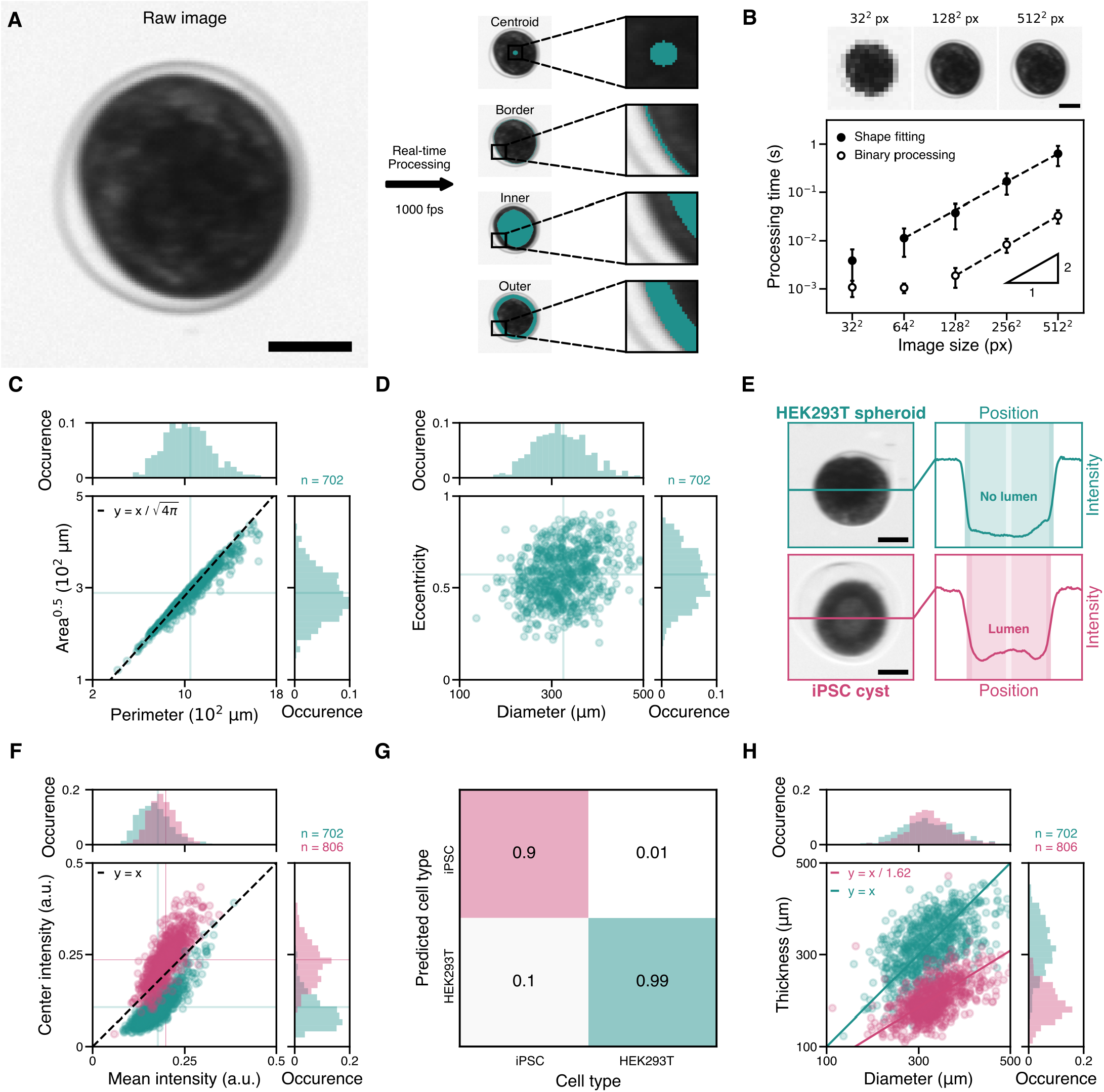
On-the-fly image processing for MCA classification. (**A**) Determination of the regions of interest in the raw image with binary mask processing. The hydrogel shell containing the cells is visible but is not considered in the analysis. Scale bar: 100 *μ*m. (**B**) Time-wise performance of the image processing algorithm. Reducing the image size down to 256 × 256 px^2^ results in ∼ 3 *μ*m*/*px and increases the processing time while maintaining satisfactory image definition. Scale bar: 100 *μ*m. (**C**) Plot of the area-to-perimeter ratio in HEK293T spheroids. The broad distribution of sizes results from variations in initial cell densities during the production of spheroids. (**D**) Plot of the eccentricities of spheroids with respect to their sizes. This refined morphological characterization allows to identify target MCAs with a given sphericity. (**E**) Transmitted intensity profiles in HEK293T spheroids and iPSC cysts. A bright center spot is observed in cysts due to the presence of a weakly absorbing lumen, whereas the increased cell density at the center of spheroids results in a minimum of intensity at the center. (**F**) *Center intensity* vs. *Inner intensity* plot for cysts (pink) and spheroids (blue). The two populations are separated by the *y* = *x* line. (**G**) Performance of the deterministic classification algorithm using the *Center intensity >Inner intensity* criterion. The data presented in the confusion matrix yields 99% precision and 90% recall scores. (**H**) Plot of the tissue thickness estimated with Beer-Lambert’s law as a function of the diameter of the MCAs. The spheroid data is used to measure a typical attenuation length *h*_0_ = 1.4 ± 0.1 × 10^2^ *μ*m, which is then used to estimate the thickness *h* of the cell monolayer in iPSC cysts. We find a linear scaling law with *h* = *D/*1.62. In (**C**), (**D**) and (**F**), solid horizontal and vertical lines indicate the sample means.

To validate this basic morphological analysis, we first measure the distribution of sizes in a population of HEK293T spheroids (*Fig. 2C*). This operation is required in toxicity assays, where MCAs of well-defined sizes must be used. We observe that the areas *A* and perimeters *P* of spheroids are correlated with a scaling law *A*^0.5^ = *k* × *P*. At the population level, we find as a first approximation *k* (4*π*)^0.5^, which is the expected value in the case of perfectly spherical objects. This legitimates the definition of an equivalent diameter *D* = 2 × (*A/π*)^0.5^, which yields a mean spheroid diameter *D* = 3.3 ± 0.6 × 10^2^ *μ*m. This heterogeneity around the mean diameter arises from the inherent variation in initial cell concentration in the alginate shells, which tends to increase over time as the number of cells grows exponentially. Apart from their sizes, spheroids may also differ in shape, as they are not necessarily spherical. They are appropriately characterized by their eccentricity *e*, defined by the relation *e*^2^ = 1 − *b*^2^*/a*^2^, where *b* and *a* are the semi-minor and semi-minor axis obtained with an ellipse fit (*Fig. 2D*). We measure a mean eccentricity *e* = 0.57, corresponding to a 17% variation between *a* and *b*. Note that *e* is an apparent eccentricity, because the image is the projection of an ellipsoid on the imaging plane, which depends on the orientation of the spheroid with respect to the camera. Here, this eccentricity mainly originates from the very shape of the hydrogel shells to which the spheroids conform upon reaching confluence. From a more general perspective, *e* may be used as a marker to study the anisotropic growth of MCAs, as observed typically in gut organoids (*36*). Overall, this morphological approach allows to measure quantitative shape-related parameters in large populations of spheroids, which is utilizable for subsequent sorting of spheroids with a target size or shape.

We then extend the methodology to a phenotype-driven analysis. Here, we distinguish spheroids from cysts through the analysis of the radial intensity profile of light transmitted through the MCAs (*Fig. 2E*). In the*Outer* region, the intensity of a cyst and a spheroid are similar because they are equally composed of cells. In contrast, their intensities in the Center region differ due to unequal cellular contents. In the case of spheroids, the length of absorbing material through which photons pass is maximum at the center. This corresponds to a minimum of transmitted light due to increased absorption and scattering. Conversely, in cysts, the deficit of absorbing material in the lumen results in a local maximum of transmitted light in the vicinity of the center.

Therefore, we discriminate cysts from spheroids by comparing the *Center intensity* to the *Inner intensity*, where the *Center intensity* is taken as the mean intensity over a small disk of 11 px in diameter (empirically chosen) around the centroid, and the *Inner intensity* is taken as the mean intensity within the *Inner* region. *Figure 2F* shows that cysts and spheroids are found, respectively, above and below the threshold line where *Center* and *Inner* intensities are equal, which validates the relevance of this discrimination criterion. Altogether, this approach yields 99% precision (true positive / true positive + false positive) and 90% recall (true positive / true positive + false negative) scores (*Fig. 2G*), which constitutes an excellent classification algorithm. These scores are slightly degraded by the fact that not all iPSC cysts exhibit a lumen: when they reach confluence in their alginate shell, an inward cell growth is observed, which fills the lumen (*37*).

Additionally, we correlate these morphological and phenotypical approaches to achieve a more comprehensive description of the MCAs. Using Beer-Lambert’s law for light attenuation in an absorbing medium, the thickness *h* of a sample is calculated from the measurement of the transmitted light intensity *I* using *h* = *h*_0_ × log(*I/I*_0_), where *h*_0_ is the attenuation length of the medium, and *I*_0_ is the incident optical intensity. Determining *h*_0_ has been shown to bear information on the chemical composition of a tissue, which makes it useful in diagnostics (*38, 39*). Here, we estimate *h*_0_ by fitting the value of log(*I/I*_0_) to the diameter *D* of HEK293T spheroids with a linear model. Although the ellipsoid nature of spheroids introduces a geometrical bias, we observe a good fit at the population level, as shown in *Fig. 2H*. We measure *h*_0_ = 1.4 ± 0.1 × 10^2^ *μ*m, which is in agreement with the typical value of 1.3 × 10^2^ *μ*m measured for kidney tissue (*40*). Though the optical properties of a tissue depend on the cell types it comprises, we also use this model to estimate the thickness of the cell monolayer of iPSC cysts with the value of *h*_0_ previously determined. At the center, assuming the attenuation in the lumen is negligible, the estimated thickness *h* should be equal to twice the thickness of a cell. We derive the scaling law *h* = *D/*1.62, which indicates that cysts with a larger diameter have a thicker cell monolayer. Remarkably, previous studies have shown that iPSC cysts exhibit an unusual anisotropic growth where the monolayer thickens as the cysts grows, which supports the validity of our results (*41*). Overall, using this combined pheno-morphological analysis in large populations of MCAs yields quantitative information on complex physiological mechanisms which could hardly be detected with the analysis of only a few samples.

### Continuous flow of MCAs and encapsulation in millimetric drops

The encapsulation of MCAs into liquid drops is achieved by flowing them into a square glass capillary using a simple polydimethylsiloxane (PDMS) millifluidic injection chip (*Fig. 3A*). The choice of a capillary with a cross section larger than twice the diameter of the MCAs (800 × 800 *μ*m^2^) mitigates the risk of clogging and flow instabilities. In contrast to most chips used to manipulate individual cells, the capillary is placed vertically to prevent sedimentation of the aggregates, which reduces the likelihood for them to stick to the inner walls of the capillary.

**Fig. 3.**
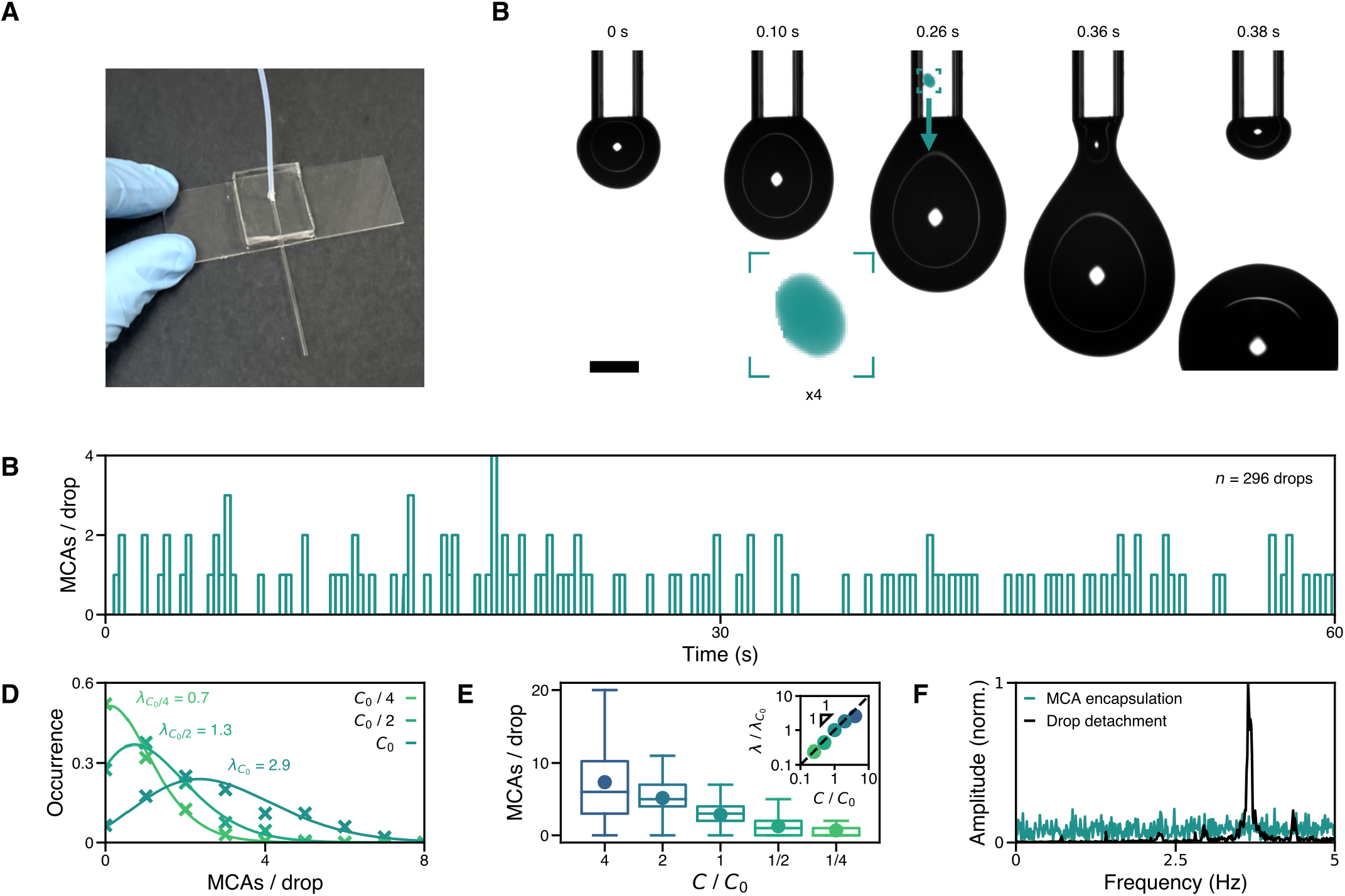
Controlled flow and encapsulation of individual MCAs into liquid drops. (**A**) Photograph of the injection microfluidic chip, with the PTFE tubing, the PDMS connection chip and the square glass capillary. (**B**) Principle of the encapsulation step. Only one MCA is imaged at a time prior to being encapsulated. The delay between imaging and encapsulation is *<* 10 ms, which ensures that each MCA is instantaneously found in the drop right after its detection. Scale bar: 1 mm. (**C**) Chronogram of the number *N*_*s*_ of spheroids per drop. Stochasticity is observed in the distribution of *N*_*s*_, as well as in the evolution of *N*_*s*_ with time. (**D**) Distributions of *N*_*s*_ for different concentrations *C* of spheroids. Data points are represented as colored crosses. Solid curves with the corresponding colors represent fits of a Poisson statistic, where the *λ* parameter is chosen as the mean *N*_*s*_ value for each distribution. (**E**) Box plot of the distributions of *N*_*s*_ as a function of *C*, normalized to a reference concentration *C*_0_ ∼ 20 MCAs/mL. Solid points represent the *λ* values. Inset: *λ* scales linearly with the dilution factor over two orders of magnitude. (**F**) Time-domain discrete Fourier transforms of the times of drop detachment and MCA encapsulation signals. The drop generation frequency distribution is peaked at 3.8 Hz while only randomness is observed in the MCA encapsulation signal.

MCAs in suspension in a culture medium solution are gently flowed through a polytetrafluoroethylene (PTFE) tubing and through the chip using a pressure pump at a constant flow rate of typically ∼ *Q* 100 − 200 *μ*L*/*s in a laminar flow regime (*Re* ∼ 10^−2^). This range of flow rates imposes a dripping regime where millimetric drops of culture medium are produced with a constant volume *V*_*d*_ = 50 *μ*L, or equivalently a constant radius *R*_*d*_ = 2.3 mm. As shown in *Fig. 3B*, each MCA arriving at the end of the capillary falls into the current pendant drop before its detachment.

Although a fraction of each drop remains attached to the capillary (typically 5% volume fraction), we could not observe a single instance of an MCA in this remaining fraction. This ensures that when an MCA is analyzed right prior its exit of the capillary, it is found in the very next drop and not in the one that follows it, which would complicate the sorting step.

A drop-based sorting system requires control over the number of objects encapsulated in each drop to minimize the probability of co-encapsulation. Here, it requires to prevent the sedimentation of MCAs in the injection vial by gently agitating it. We characterize this encapsulation step by measuring the distribution of *N*_*s*_, the number of MCAs found in each drop (*Fig. 3C* and *Movie S1*). At a reference concentration of *C*_0_ ∼ 1000 MCAs in 50 mL of culture medium, *N*_*s*_ randomly varies between 0 and 9 MCAs per drop, independent of the contents of the preceding drop. In a homogeneous suspension, *N*_*s*_ is expected to follow a Poisson statistic. The probability *P* of finding *N*_*s*_ aggregates in a drop is therefore given by *P* (*N*_*s*_) = exp^−*λ*^ *λ*^*N*^*s /N*_*s*_! where *λ* = *C* × *V*_*d*_ is the average number of objects per drop (*42*). In *Fig. 3C*, we vary the initial concentration *C* of MCAs and plot the corresponding distributions of *N*_*s*_. We observe an excellent agreement of the experimental data with theoretical Poisson distributions. Though this statistic does not provide deterministic control over the number of MCAs in each drop, we use it to estimate the number of coencapsulation events. For instance, setting *λ* = 0.1 results in 90% empty drops and *<* 0.5% co-encapsulations, which leaves ∼ 10% drops containing a single object (35). However, with *V*_*d*_ = 50 *μ*L, this configuration imposes a concentration *C* = 2 MCAs/mL, which would reduce the sorting throughput and waste large amounts of culture medium. We instead opt for a typical concentration of 5 MCAs/mL to reduce the processed volume and increase the throughput up to 1 MCA/s. The residual co-encapsulations cases (∼ 10%) are mitigated by not sorting the corresponding drops, as detailed in a following section.

Additionally, we emphasize that conveying MCAs within the capillary has no effect on the regularity of the drop production rate. *Figure 3F* shows the relative amplitudes of the discrete Fourier transforms of the drop pinch-off timestamps and of the times of arrival of MCAs at the exit of the capillary. While MCAs arrive at random timestamps, drops are generated at a constant frequency of 3.8Hz, independently of the number of MCAs encapsulated. This regularity legitimates the choice of a fixed delay time between the detection of a target MCA in the capillary and the actuation of a trigger signal to deflect the drops.

### Acoustic-driven actuation of millimetric liquid drops

In conventional flow cytometers, drops in the *R*_*d*_ ∼ 100 *μ*m radius range are actuated using the electrostatic force by applying a constant electric field perpendicular to the freefall trajectory of the drops (*25*). However, since this force scales as 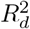, the electric field should be increased by more than 2 orders of magnitude for millimetric drops, typically reaching over 10^5^ V.m^−1^. This would lead to the breakup of the drops (*43*). On the contrary, placing a millimetric liquid drop in a standing-wave acoustic field yields an ARF strong enough to manipulate the drop on a surface (*44*), and even to keep it in acoustic levitation (*45*). Physically, a standing-wave acoustic field is scattered by the presence of a liquid drop, which creates second-order pressure and velocity fields. These secondary fields have non-null time averages and thereby contribute to a net force *F*_*a*_ on the surface of the drop, parallel to the direction of propagation of the incident acoustic waves (*46, 47*). When *R*_*d*_ is small relatively to the acoustic wavelength *λ*_*a*_, this acoustic force writes:

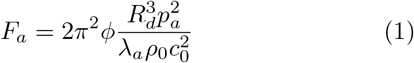

where *p*_*a*_ is the acoustic pressure amplitude, *ρ*_0_ is the density of air, *c*_0_ is the speed of sound in air and *ϕ* ∼ 5*/*6 is the acoustic contrast factor between air and water (*48*). *F*_*a*_ is directed from the pressure antinodes towards the pressure nodes. This model yields a good approximation as long as *R*_*d*_*/λ*_*a*_ *<* 0.3 (*49*). The scaling of *F*_*a*_ with the volume of the drop is what makes it suitable for the manipulation of large drops against gravity.

To deflect falling drops at the exit of the encapsulation capillary, we generate a standing-wave pressure field along the *y* axis with two hemispherical arrays of 36 ultrasonic transducers in phase operating at 40 kHz (*Fig. 1*). This sets *λ*_*a*_ = 8.6 mm with *R*_*d*_*/λ*_*a*_ = 0.25, and the resulting standing-wave acoustic field has an inter-node distance *λ*_*a*_*/*2 = 4.3mm. In this configuration, it exhibits a plane symmetry with respect to the *xy* plane located at its center in the *y* direction, which corresponds to a pressure antinode (*50*). For this reason, the exit of the capillary needs to be placed slightly off-center, typically *δy* = *λ*_*a*_*/*8, so that the drops fall between a node and an antinode where the ARF is stronger. *Figure 4A* shows the deflection of a pendant drop upon activation of the acoustic field during its detachment. This occurs in three phases. First, the pendant drop remains attached to the capillary under the action of surface tension force *F*_*γ*_. As it grows due to the constant flow of fluid through the capillary, it reaches a critical radius 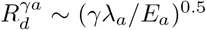 given by the *F*_*γ*_*/F*_*a*_ balance. Above 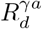, the ARF overcomes the capillary forces in the *y* direction, which deflects the drop horizontally. After reaching a second critical radius 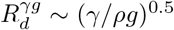 given by the *F*_*γ*_*/F*_*g*_ balance where *ρ* = 10^3^ kg.m^−3^ is the density of water and *g* the gravitational acceleration constant, the pendant drop eventually detaches from the capillary with a net momentum along the *y* direction.

**Figure 4.**
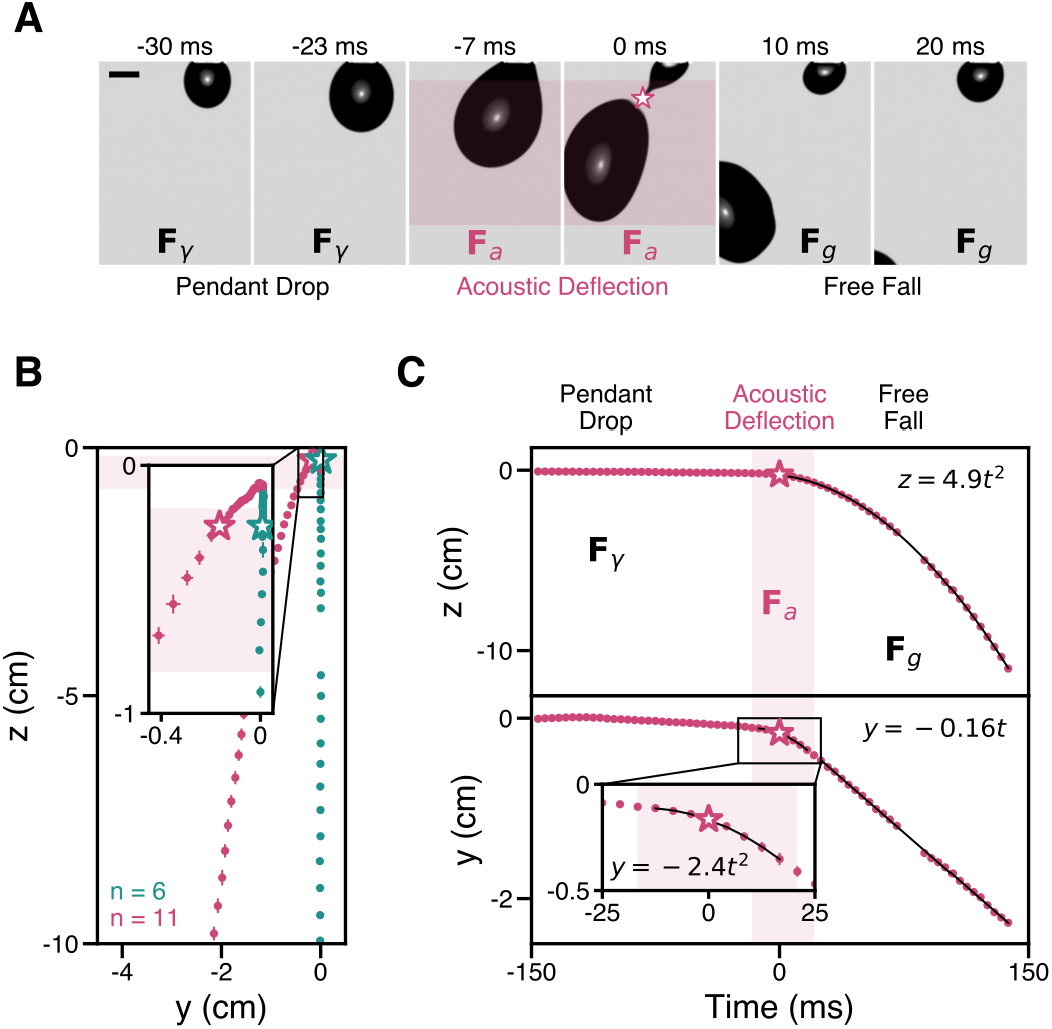
Deflection of pendant drops using the ARF. (**A**) Images of the behavior of a pendant drop in a standing-wave acoustic field. While *R*_*d*_ is smaller than a critical radius 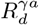 (for *t <* −23 ms), surface tension dominates, and a pendant drop is observed. Around the detachment point (for −7 ms *< t <* 0 ms), the ARF becomes prominent and the drop is horizontally deflected. Once *R*_*d*_ becomes greater than the capillary length 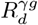, the drops detaches (pink star at *t* = 0 ms). It then quickly reaches its terminal velocity and escapes the acoustic field in a free fall where it is only submitted to its weight *F*_*g*_. Scale bar: 1 mm. (**B**) Trajectories of the centroid of falling drops. Deflected drops deviate from the *z* axis with a typical angle ∼ 40 deg at the exit of the capillary, which allows to collect them in a separate vial. Points are spaced 4.2 ms apart. (**C**) Vertical and horizontal positions of drops’ centroids as a function of time. We estimate the intensity *F*_*a*_ of the ARF using Newton’s second law of motion, yielding *F*_*a*_ = *mv*_0_*/*Δ*t*_*a*_ = 2.4 × 10^−4^ N where *m* is the mass of a drop, *v*_0_ its terminal horizontal velocity and Δ*t*_*a*_ its time of flight in the acoustic field. In (**B**) and (**C**), dots represent the mean positions and bars represent the *y* and *z* standard deviations.

To compare the dynamics of deflected and non-deflected drops along their fall, we analyze the trajectories of their centroids (*Fig. 4B* and *Movie S2*). We first observe that the deflection of the drops is initiated before their detachment with excellent reproducibility. When operating at a rate of 2 drops/s, the ARF imposes a lateral deflection Δ*y* = 1 cm in about Δ*z* = 3 cm falling distance. This is sufficient to collect deflected drops and non-deflected drops in separate vials. We successfully separate individual drops at flow rates up to 500 *μ*L*/*s, or equivalently 10 drops*/*s, which corresponds to the dripping/jetting transition.

A more detailed analysis of the trajectories provides information on the intensity of the ARF acting on large, deformable liquid drops, which could otherwise only be calculated at the cost of subtle corrections that are beyond the scope of the present work. We first track the coordinates of the drops’ centroids over time to obtain the vertical and horizontal displacements *z*(*t*) and *y*(*t*) (*Fig. 4C*). Strictly speaking, *F*_*a*_ is the component of the ARF along the axial *y* axis, but the ARF also has components in the radial *x* and *z* directions (*48*). However, we neglect them in a first order approximation as they are reportedly ∼ 5 times weaker than the axial component *F*_*a*_ (*49*). In our system, the *x* component is cancelled out anyway since the capillary is placed in the *x* = 0 plane of symmetry. We make the bold simplification that *F*_*a*_ is approximately constant in space and time along the drop’s trajectory in the acoustic field, indicated by the shaded region in *Fig. 4C*. Under these assumptions, the initial speed at the exit of this zone of actuation is simply *v*_0_ = *F*_*a*_Δ*t*_*a*_*/m*, where Δ*t*_*a*_ is the time of flight of the drop in the acoustic field and *m* = 5 × 10^−5^ kg is the mass of the drop. In the absence of friction, the drop is in free fall after exiting the acoustic field and *z*(*t*) = *gt*^2^*/*2. *Figure 4C* shows an excellent agreement of our experimental data with this simple kinetic model, which legitimates the omission of the *z* component of the ARF. From the fit of *y*(*t*), we obtain *F*_*a*_ = 2.4 × 10^−4^ N ∼ *mg/*2. This force would be too weak to quantitatively deflect a falling drop that has already reached its terminal velocity, but here a strong initial deflection of ∼ 40° is observed (*Fig. 4B*) because it occurs while the drop has little to no vertical speed. Using *Eq. 1* to estimate the acoustic pressure in the region of deflection, we obtain *p*_*a*_ = 1.2 × 10^3^ Pa = 1.6 × 10^2^ dB, which is on par with values previously reported for similar acoustic systems (*45, 49–51*). Note that due to the high contrast in acoustic impedance between air and water, the water drops are nearly impenetrable to acoustic waves. The air-water interface thus protects the encapsulated MCAs from damage by the relatively intense acoustic field.

### Sorting large populations (n *>* 1000) of MCAs in a continuous flow

After optimization of the detection, encapsulation and actuation modules, efficient sorting *simply* requires the synchronization of these steps. Here, the suspension of MCAs is flowed at constant rate, which sets a drop generation frequency of 3.8 drops*/*s with a period *T*_*d*_ = 0.26 s between successive drops. Since the images are acquired at ∼ 1 mm above the capillary exit where the MCAs flow at ∼ 10 cm.s^−1^, an MCA is found in the drop ∼ 0.01 s after being detected. Therefore, upon detection of a target MCA, only the very next drop must be sorted. To do so, we activate the acoustic field for a duration of 1.5 × *T*_*d*_, right after the image processing returns a positive result. Additionally, we implement a simple decision tree to minimize potential co-encapsulation events that could deteriorate the sorting efficiency (*Fig. 5A*). When two objects are detected within a duration Δ*t < T*_*d*_, the next drop is not deflected if one or more of the objects is identified as non-target.

**Fig. 5.**
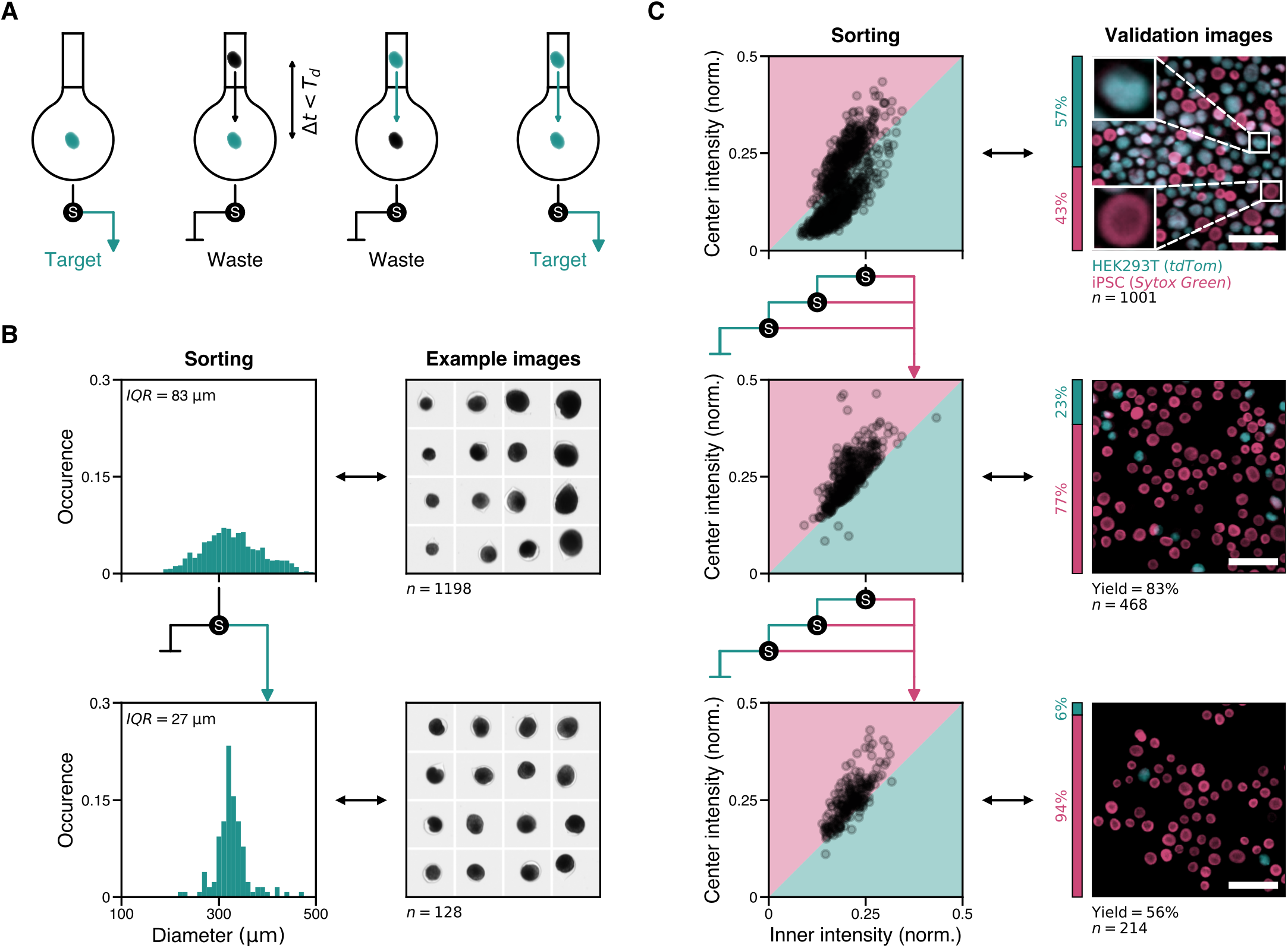
Sorting performance of ImOCAS. (**A**) Illustration of the co-encapsulation mitigation strategy. Drops containing at least one non-target object are discarded: if two objects or more are detected within Δ*t < T*_*d*_ and one of them is classified as waste, the following drop is not sorted. (**B**) Histograms of distributions of sizes and typical images of HEK293T spheroids before and after sorting them using two threshold radii *D*_*min*_ = 320 *μ*m and *D*_*max*_ = 336 *μ*m. The radius distribution is initially broad before sorting (top row, *IQR* = 83 *μ*m) and is narrowed down after sorting (bottom row, *IQR* = 27 *μ*m). The median diameter *M*_*D*_ = 329 *μ*m remains unchanged. Not all spheroids fit in the [*D*_*min*_, *D*_*max*_] range because the definition of *D* depends on the orientation of a spheroid with respect to the camera, hence highly non-spherical spheroids are imperfectly characterized by the measure of *D*. Images are 800 × 800 *μ*m^2^. (**C**) Sorting strategy and validation fluorescence images for the sorting of iPSC cysts from a sample where they are mixed with HEK293T spheroids. We use the *Center intensity > Inner intensity* criterion to detect target cysts, as shown by the shaded regions in the left column plots. The fluorescence images shown in the right column are acquired afterwards to measure the purity *P* and yield *Y* sorting metrics: *P* = 77% and *Y* = 83% after one sorting step, *P* = 94% and *Y* = 56% after two sorting steps. Scale bars: 1 mm. The *S* disks represent one sorting step and the arrows indicate the sorting procedure.

One key challenge in large-scale biological assays is ensuring homogeneity within the sample population. For instance, to evaluate the impact of a drug in a population of spheroids, their initial size distribution must be uniform. Thus, we first evaluate the efficiency of ImOCAS by sorting spheroids based on their size. We compare the diameter *D* of a spheroid measured in an image with two threshold radii *D*_*min*_ and *D*_*max*_ set by the user. Due to the non-spherical shape of a spheroid, the definition of its radius depends on its orientation with respect to the imaging plane, which introduces variability in the determined value. The objective of this sorting step is to reduce the width of the size distribution to obtain a homogeneous sample, rather than to accurately select a size range. As shown in *Fig. 5B*, we first use ImOCAS in a cytometry mode to characterize our starting population of *n* = 1198 spheroids with a median value *D*_*M*_ = 329 *μ*m, and an interquartile range *IQR* = 83 mum. The whole distribution of *D* extends between 117 − 558 *μ*m. We define two thresholds *D*_*min*_ = 320 *μ*m and *D*_*max*_ = 336 *μ*m that encompass 10% of the spheroids centered around the median diameter. We then use ImOCAS to sort the spheroid population accordingly, and we finally use it to screen the resulting sample. After sorting, the width of the size distribution is considerably reduced, with *IQR* = 27 *μ*m, without modifying the median diameter *D*_*M*_. This demonstrates the efficiency of ImOCAS to sort spheroids based the measurement of a continuous morphological parameter (here, their diameters).

ImOCAS is also able to sort MCAs based on their phenotypical signature, which finds applications in organoid screening. For instance, in the case of lumenized iPSC cysts, live samples clearly exhibit a lumen while differentiated tissues resemble plain spheroids (*37*). Here, we mimic this situation using a heterogeneous sample containing *N*_*T*_ = 432 cysts (Target) stained with Sytox Green and *N*_*W*_ = 569 spheroids (Waste) stained with tdTom. The sorting is again performed based on bright-field image analysis, but we use fluorescent dyes to subsequently validate the sorting results via epifluorescence imaging. We use the *Center intensity > Inner intensity* criterion as a signature of the presence of a lumen to sort the two sub-populations (*Fig. 5C*). Since drops containing two or more MCAs are not sorted, we compensate for the expected decrease in yield with a multiple-step sorting approach. After sorting the initial sample population, the contents of the waste vial are re-sorted to recover the discarded target objects, and finally the contents of the resulting waste vial are also sorted. To be selected, an MCA therefore needs to be marked once as a target among three sorting steps (*Fig. 5C*, rows 1 and 2). We assess the sorting quality by measuring the purity *P* = *N*_*T*_ */N*_*W*_ and the yield 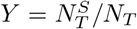, where 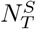 is the number of sorted targets. Starting from *P* = 43%, this first sorting step removes 75% of the spheroids while keeping *Y* = 83% of the cysts to obtain *P* = 77%. The discrepancy between this satisfactory purity rate and the excellent precision of the image processing algorithm mainly emerges from the aggregation of some MCAs into clusters of two or more objects, which cannot be properly classified. However, since the whole sorting procedure is short (*<* 1 h), the purity is easily further increased with a second sorting step leading to *P* = 94% (*Fig. 5C*, bottom row).

Overall, ImOCAS is suitable for sorting large populations of MCAs based on both morphological and phenotypical parameters at a high throughput, typically 100 times faster than manual sorting.

## Discussion

The use of 3D cell models in both fundamental biology and biomedical applications has long been hampered by the absence of robust protocols to produce and analyze these complex objects. In the last 20 years, numerous methods based on microfluidics and microfabrication techniques have emerged and now allow the reproducible, high-throughput production of MCAs. However, to perform statistically relevant biological assays on a homogeneous population of MCAs, one also needs to sort them to reduce the inherent morphologic and phenotypic variability of these biological samples. There is therefore a growing need to implement strategies for analyzing and sorting large populations of MCAs. Counter-intuitively, simply scaling up the microfluidic devices that were designed to manipulate single cells is highly challenging, mainly due to the prominence of inertia and gravity at the scale of MCAs (50 *μ*m − 5 mm).

To the best of our knowledge, flow-based organoid and spheroid sorting using image analysis has only been achieved by a handful of commercial systems (*52, 53*). However, these systems are limited by the impossibility to tailor them for custom applications and by their dependence only on pre-defined morphological parameters or on averaged intensity levels rather than comprehensive image analysis.

To address these limitations, we developed ImOCAS, a drop millifluidic system which performs automated screening and sorting experiments on large populations of MCAs. Like a traditional flow cytometer, ImOCAS operates in three successive steps: detection of a feature in MCAs in flow, encapsulation of single MCAs in liquid drops and actuation of the drops of interest. Here, each step is sought to be adapted to the manipulation of MCAs instead of single cells. First, the morphological and phenotypical signatures of MCAs are detected using rapid image processing while the MCAs flow inside a glass capillary. Then, individual MCAs are encapsulated in millimetric drops of culture medium in a dripping mode. Finally, the drops are actuated using the ARF upon the generation of a standing-wave acoustic field in the vicinity of the drops.

The main strengths of ImOCAS are its versatility regarding the detection parameters and its potential to be used in any biology laboratory since it does not require expensive nor complex apparatus other than a computer with standard performance for image analysis and a fluid control system. Its small size (40 × 20 × 20 cm^3^) makes it fit inside a biosafety hood if working in a sterile environment is required. The choice of a label-free imaging method also makes it compatible with live samples.

The current sorting throughput of 1 Hz may not be compared to the kHz rates reported in single-cell sorting due to the 1000-fold volume ratio between the sorted objects. In ImOCAS, the sorting throughput is only limited by the frequency of drop production, which could be increased by reducing the size of the drops. This may be achieved using a piezo-electric actuator to vibrate the capillary and thereby trigger the detachment of the drops, for instance. Further optimizing the flow upstream of the capillary to increase the fraction of non-empty drops constitutes a promising approach, which was already observed with the encapsulation of smaller particles in microfluidic chips (*54*). On the detection side, although ImOCAS already allows a thorough pheno-morphological analysis of MCAs, modifications could be made to meet specific requirements. For instance, the use of a 3D imaging method would offer a more comprehensive representation of the structural complexity of MCAs. However, this poses technical challenges as ImOCAS requires usage of a long working distance objective in order not to interfere with the acoustic field in the deflection zone. Regarding the actuation step, there is no physical limitations to the use of more complex acoustic fields to sort drops in more than two categories. In particular, the acoustic nodes may be accurately re-positioned in real time with a simple phase shift between the two arrays of transducer, hence changing the direction of deviation along the *y* axis. Overall, ImOCAS is here presented in its most general form and could be easily tailored for custom applications.

ImOCAS may rapidly be adopted in biology facilities as a robust approach to analyze and sort complex 3D biological models. Thanks to its versatility, it may be applied broadly to standardize drug testing experiments in MCAs by first ensuring the homogeneity of the tested MCAs and then screen the results. It may also be used to detect rare events that would be left unnoticed when analyzing only a small number of MCAs. We anticipate that the compatibility of ImOCAS with virtually any organoid and spheroid samples will unveil new insights in 3D biology, in both fundamental and applied research.

## Materials and Methods

### Cell culture

#### 2D culture

HEK293T cells obtained from the Bordeaux Institute of Oncology were infected with a lentivector backbone expressing tdTom with Nter Palmitoylation (under the control of hPGK), using standard methods. Infected cells constitutively express the tdTom fluorescent molecule at their membrane. Cells were then cultured in culture-treated plastic flasks (Corning, cat. no. 353136) and maintained in Dulbecco’s Modified Eagle Medium (DMEM) (Biowest, cat. no. L0103-500) supplemented with 10% fetal bovine serum (Capricorn, cat. no. FBS-16A), 5% penicillin-streptomycin (Gibco, cat. no. 15140122). iPSC cells purchased from Corriell Institute (AICS-0023) were cultured in culture-treated plastic flasks (Corning, cat. no. 353107) coated with 5% Matrigel (Corning, cat. no. 354234), maintained in mTeSR Plus (StemCell, cat. no. 100-0276). Both cell types were maintained under water-saturated 5% CO2 atmosphere at 37°C.

#### Formation of MCAs

Cell aggregates were formed using the CCT method, previously described in (*16*). Briefly, cells were detached from their flasks using trypsin (Biowest, cat. no. L0930-100) for HEK293Ts and Accutase (Stem-Cell, cat. no. 07920) for iPSCs. Cells were resuspended in their appropriate culture medium and mixed with Matrigel (Corning, cat. no. 356234) at 50% volume fraction. The resulting solution was injected in a 3D printed microfluidic chip together with a solution of sorbitol 300 mM (Sigma-Aldrich, cat. no. S1876) and a solution of 2% alginate (AGI, cat. no. I3G80) to form a composite liquid jet consisting in a triple co-flow of the cells/Matrigel mix, sorbitol and alginate solutions. The jet fragmentation resulted in the formation of alginate shells, collected in a calcium bath to trigger alginate reticulation, hence forming solid core-shell structures with cells embedded in Matrigel in the core. The seeding density ranged from 10 − 100 HEK293T cells per shell, and 1 − 10 iPSC cells per shell. In the case of iPSCs, ROCK inhibitor (Tocris, cat. no. 1254/10) at 10 mM was added and replenished at 1*/*1000 dilution every day for the first 48 h after formation of the shells.

Since alginate is optically transparent, it does not alter the image-based analysis of the aggregates, and we effectively probe only the morphology and size and the MCAs contained inside the alginate shells. Any other classical method to produce MCAs may be used, providing that the aggregates can be resuspended in a solution. 3D culture Cell-laden alginate shells were cultured in the appropriate culture medium for 7 days, with medium replacement every 2 − 3 days. The cells were then fixed in a solution of 4% paraformaldehyde (Sigma-Aldrich, cat. no. 47608) overnight and stored in glucose free and phenol red free DMEM (Gibco, cat. no. A1443001) at 4°C. The nuclei of iPSC cells were stained with SYTOX Green after fixation (NucGreen Dead 488, Thermo Fisher).

### Detection

#### Image acquisition

In the glass capillary, cell aggregates were illuminated with a white light LED (MWWHL4, Thorlabs), collimated with a single convergent lens (AC254-050-A-ML, Thorlabs). The transmitted light was collected through a zoom lens (1-60135 6.5X, Navitar) and acquired with a CMOS monochromatic camera (acA720-520um, Basler) at ∼ 100 fps. The typical image size was 256 × 256 px^2^ for a field of view of 800 × 800 *μ*m^2^, yielding ∼ 3*μ*m*/*px (i.e., 6 *μ*m resolution).

#### Image processing

On-the-fly image processing was conducted using a Python script with the Numpy, OpenCV and Scipy libraries. First, the intensity of each pixel was normalized against that of an image without objects. Subsequently, an intensity threshold was applied to identify cell aggregates, represented by pixels with lower intensity. The centroid of the identified region was calculated, and a 11-px radius disk was drawn around it to define the *Center* region. The outermost border of the identified region constituted the *Border* region. By eroding the region by 6 px, we obtained the *Inner* region, and the difference between the initial region and the *Inner* region yielded the *Outer* region. Mean intensities across each region were calculated by summing px intensities and dividing by the area of the region.

### Encapsulation

#### Millifluidic injection chip

Cell aggregates were flowed from a standard Falcon tube (catalog no. 352098, Corning) into an 800 × 800 *μ*m^2^ square glass capillary (VitroCom, cat. no. 8280) through a PTFE tubing with 1.06 mm internal diameter (Fisher Scientific, cat. no. 11949445). The connection between the tubing and the capillary was made using PDMS (Sylgard 184, Dow Corning). A mold was fabricated by sticking a capillary onto a glass slide, then pouring the PDMS on top of it before baking it at 60°C for 4 h.

#### Fluid control system

The aggregate suspension was transferred into the encapsulation chip using a pressure controller (MFCS-EZ, Fluigent). The driving pressure was set to ∼ 40 bar to achieve flow rates of 100 − 500 *μ*L.s^−1^, or 2 − 4drops*/*s, using a PTFE tubing of ∼ 30 cm in length. This setup maintained a steady flow rate without requirement for a flow meter.

#### Sample stirring

Due to their size, organoids and spheroids in suspension tend to sediment at the bottom of a 50 mL vial within ∼ 10 s. It is therefore necessary to mix the suspension continuously. However, using a magnetic stirrer would harm the aggregates. Instead, the vials containing the suspension were mounted onto a servo motor (MG996R, Tiankon-gRC), controlled by a microcontroller (Uno Rev2, Arduino), to perform back-and-forth rotations with an amplitude of approximately 100°.

### Actuation

#### Acoustic field design

Our acoustic deflection system draws inspiration from prior research on acoustic levitation setups (*45*). The system comprises two spherical caps designed to hold ultrasonic transducers. These caps were 3D printed using a Digital Light Processing 3D printer (D4K Pro, Envisiontec) and photocurable resin (HTM 140 V2, Envisiontec). The radius of curvature of the caps was set to 86 mm, or equivalently to 10 acoustic wavelengths, ensuring an effective focusing effect while maintaining sufficient space to manipulate the capillary between the two caps. Each cap accommodates 36 ultrasonic transducers (MA40S4S, Murata), arranged along concentric rings and electrically connected in parallel. A 40 kHz square signal, generated by a waveform generator (T3AFG30, Teledyne Lecroy), was amplified with an L298N motor driver electronic chip powered at 9.5 V before being transmitted through the transducer circuit. The two spherical caps were aligned to ensure that each transducer on one cap is facing its counterpart on the other, hence generating a standing-wave acoustic field with spherical iso-phase planes.

#### Positioning the acoustic field relatively to the capillary

The spherical focus of the acoustic field resulted in a maximum pressure amplitude at the system’s center. The intensity of the resulting ARF had local maxima located halfway between the nodes and the antinodes of the standing pressure field, which had an inter-node distance *λ*_*a*_*/*2 = 4.3 mm. Therefore, the tip of the glass capillary had to be positioned along the axis of symmetry of the acoustic field, but slightly off-center (typically *λ*_*a*_*/*8) to maximize the local pressure gradient. In practice, the optic and fluidic apparatus were kept still, and the spherical caps were moved along the *y* axis until satisfactory drop deflection was observed, and along the *x* axis until no deflection was observed in this direction.

## Acknowledgments

We thank L. Andrique for identifying the requirements and potential applications of the development of a system to sort spheroids and organoids. We thank D. Baresch and M. Beaudoin for fruitful discussions regarding the acoustic setup. We also thank D. Baresch for critical reading of the manuscript. We acknowledge the assistance of X. Bril and A. Tizon with the electronic setup.

## Author contributions

LR, PN and JCB conceptualized the project. LR and TB conceived the fluidic setup. LR, AmB and LB conceived the optical setup. CD, AJ, AdB and LH produced the biological samples. LR developed the image analysis scripts and models, designed the acoustic setup, build the sorting platform and performed the experiments. PN and JCB supervised the project. LR, PN and JCB wrote the manuscript.

## Data and materials availability

All data needed to evaluate the results presented here are present in the main text and/or in the *Supplementary Materials*. The analysis codes used in this study can be provided by the corresponding authors upon request.

