## Supplementary Materials for "Pheno-morphological screening and acoustic sorting of 3D multicellular aggregates using drop millifluidics"

**Movie S1**
Encapsulation of multicellular aggregates (MCAs) into drops of culture medium at the exit of a glass capillary. The concentration is $C \sim 5$ MCAs/mL, which results in $N_s = 0.7$ MCAs/drop on average, and 10% of drops containing 2 MCAs or more. Speed slow down four times (100 fps).

**Movie S2**
Deflection of water drops using the Acoustic Radiation Force (ARF) for sorting purposes. The injection glass capillary is placed at the vicinity of the center of symmetry of two hemispherical arrays of ultrasonic transducers operating at 40 kHz. While the transducers are switched off, drops free fall vertically into a collection vial. Upon activation of the transducers, a standing acoustic field is generated, which results in the application of an ARF at the surface of the drop while they still attached to the capillary. After they detach from it, they enter a free falling regime, but with an initial horizontal speed which allows to collect them in a separate vial. Actual speed (120 fps).
